# Light propofol anaesthesia for non-invasive auditory EEG recording in unrestrained non-human primates

**DOI:** 10.1101/2025.03.24.644890

**Authors:** T. Piette, C. Lacaux, M. Scheltienne, V. Sterpenich, M. Isnardon, V. Moulin, A. Cermolacce, D. Grandjean, A. Meguerditchian, E.C Déaux, A-L. Giraud

## Abstract

Non-invasive electroencephalographic (EEG) experiments have been instrumental in advancing our understanding of the brain mechanisms involved in the production and perception of sounds and human speech. Performing similar experiments in non-human primates (NHPs) would help further deepen our knowledge by allowing us to investigate the evolutionary roots of these processes. However, performing EEG on NHPs is a challenge, given its sensitivity to motion artefacts, device cost and durability, and animal training requirements. For these reasons, most attempts have used invasive intracranial recordings, which led us to develop an alternative that minimises stress and prioritises animal welfare. By using mild propofol sedation, neurophysiological experimentation can easily be integrated into the routine sanitary checks of captive animals and allows the optimisation of both EEG quality and animal welfare. To assess the influence of propofol on brain activity in NHPs, we sedated three olive baboons (*Papio anubis*), scored their sleep stages under different doses, and recorded auditory event-related potentials (ERP)in response to grunts. Analyses of the EEG recordings with regards to sleep stage and ERP components indicate that at low dose (< 0.1mg/kg/h), propofol induces a light sleep state conducive to recording stimulus-elicited auditory activity. Overall, this experiment confirms the use of propofol sedation as an appropriate technique to study auditory processes through unrestrained, non-invasive EEG in NHPs.

## INTRODUCTION

Over the past decades, neuroimaging technologies have significantly advanced language research, providing critical insights into acoustic and speech processing. Among these technologies, electroencephalography (EEG) has played a central role in uncovering key mechanisms underlying human communication, such as the organisational role of brain oscillations, and has been instrumental in language disorder research and hearing aid development ^1–3^. EEG can also be used to study the evolution of language and acoustic communication, in the context of comparative studies, when equipped on non-human primates (NHPs)^4–8^. However, it presents significant challenges; EEG is highly sensitive to motion artifacts^9^, and the overall fragility of the recording equipment further complicates data acquisition in moving animals.

To circumvent these limitations, past studies have often relied on invasive intracranial recordings or used extensive training of a few individuals over several years ^4,10,11^. These studies frequently rely on primate chairs, a contention set-up that places the animals in stressful working conditions^12^. While such set-ups minimise motion artefacts and prevent damage to the devices, they greatly limit the scope and generalizability of the experiments and raise serious ethical questions.

An alternative approach for experiments that do not explicitly require animal cooperation involves the use of light anaesthesia to inhibit movement in test subjects, enabling high-quality data acquisition. Importantly, this method eliminates the need for physical restraints, reducing stress and fear in the animals. Additionally, by removing the requirement for animal training and ensuring device protection, this approach could lower costs, allowing for larger sample sizes and ultimately contributing to more robust scientific results. Yet, the major drawback to using anaesthetic drugs is the risk of inducing deep sleep stages and impairing auditory processing. Indeed, common veterinary anaesthetic procedures include injecting intramuscular ketamine combined with medetomidine or inhaled sevoflurane. However, these drugs have been shown to reduce auditory evoked potential responses and diminish oddball mismatch negativity, making them suboptimal candidates for neuroacoustic studies ^13–16^.

However, while veterinary practices have favoured these anaesthetics for their reliability in various settings, other agents may be better suited for sensory processing experiments. Among them, propofol, an intravenous drug from the alkylphenol family, shows promising potential. Unlike ketamine and sevoflurane, which significantly inhibit early brainstem auditory response in humans, propofol, when administered at low dosage, has minimal impact on auditory response latency and negligible effects on response amplitude ^17,18^. Furthermore, while propofol induces widespread slow oscillations (0.5-1H) and increased alpha oscillations (8-12Hz) in the frontal cortex^19,20^, no direct effect on other oscillations their coupling in the auditory cortex has been reported. Information on propofol’s effect on late auditory response, and especially ERP components N100 and P200 is however lacking. To date, only one study used propofol in a near-infrared spectroscopy experiment, and reported similarities in the processing of conspecific emotional vocalisations between humans and sedated baboons, highlighting propofol’s potential^21^. Importantly, propofol’s rapid anaesthetic response allows for fine-tuning anaesthesia at very low dosages^22^, enabling the induction of light sleep stages (NREM1–NREM2), where auditory processing is maintained, while avoiding deeper stages (NREM3-REM) that would impair auditory processing^23,24^.

In this study, we investigated the impact of propofol anaesthesia on auditory processing using non-invasive EEG recordings in olive baboons (*Papio anubis*). Initial pilot tests were conducted on two subjects anaesthetised with propofol at 0.2 mg/kg/h. These tests revealed conflicting results in sleep depth and auditory ERP responses between the two subjects, suggesting significant individual variability in the effects of propofol. To address these discrepancies, we performed a more detailed experiment on a third subject. Using continuous EEG monitoring, we assessed sleep levels under varying doses of propofol and compared auditory ERP responses during sevoflurane anaesthesia and the lowest tested dose of propofol (0.1 mg/kg/h). Our findings demonstrated that at low dosage, propofol (0.1 mg/kg/h) successfully induced a light sleep stage while preserving basic auditory processing capabilities, as evidenced by the presence of classical ERP components such as N100 and P200. These results confirm the potential of using low-dose propofol anaesthesia as a valuable tool for non-invasive brain recording of auditory processing in non-human primates.

## METHOD

### Subjects

The study included three healthy female baboons of respectively 12, 16, and 6 years old (Table 1). Male baboons were excluded due to their larger, thicker masticatory muscles over the temporal cortex, which could interfere with the collection of neural activity. The three individuals are part of the Primatology Station (Marseille, France). The baboons were born in captivity and housed in social groups in enclosures with access to outdoor and indoor areas. Their enclosures include climbing structures of wood and metal, along with substrate favouring natural foraging behaviours. Water is provided *ad libitum*, and the baboons receive daily feeds of monkey pellets, seeds, fresh fruits, and vegetables. For each of the three females, health assessments and daily behavioural monitoring by veterinary and animal welfare staff confirmed that they had normal hearing and no structural neurological impairments. This was further validated by T1-weighted anatomical brain imaging acquired under anaesthesia using a 3-Tesla MRI, performed as part of another ongoing research^25^. All procedures received approval from the “C2EA-71 Ethical Committee of Neurosciences” (INT Marseille) and complied with French law, CNRS guidelines, and European Union regulations.

**Table 1.** Summary information for each female baboon. Name, age, weight, anaesthetic type, and dosage for each subject, associated with details (type and duration) of the experimental protocol.

| Subject name | Age (years) | Weight (kg) | Anaesthetic | Dosage | Experiment type | Recording time (s) |
| --- | --- | --- | --- | --- | --- | --- |
| Fidji | 12 | 16.75 | Propofol | 0.20 mg/kg/h | ERP | 120 |
| Chet | 16 | 18.50 | Propofol | 0.20 mg/kg/h | ERP | 120 |
| Ozone | 6 | 13.35 | Sevoflurane | 3% | ERP | 240 |
| Ozone | 6 | 13.35 | Propofol | 0.15 mg/kg/h | Rest | 400 |
| Ozone | 6 | 13.35 | Propofol | 0.10 mg/kg/h | ERP | 240 |
| Ozone | 6 | 13.35 | Propofol | 0.10 mg/kg/h | Rhythm | 1860 |
| Ozone | 6 | 13.35 | Propofol | 0.20 mg/kg/h | Rest | 120 |
| Ozone | 6 | 13.35 | Propofol | 0.25 mg/kg/h | Rest | 120 |

### Stimuli

#### ERP

Auditory stimuli for the ERP experiment consisted of a sequence featuring a 100 ms grunt from an olive baboon unknown from our subjects, repeated either 50 or 100 times. Each grunt was followed by a randomly timed silence interval lasting between 1 and 2 seconds.

#### Rhythm

Auditory stimuli for the rhythm experiment (used in another study) consisted of a recording made of 5-second long sequences of human speech, baboon grunts, and white noise, repeated either 50 or 100 times each. Each sequence was followed by a randomly timed silence interval lasting between 1 and 2 seconds.

### General anaesthesia protocol

For all three subjects, we started with the general anaesthesia protocol routinely as part of general health assessments. First, subjects were separated from their social group and anaesthetised with an intramuscular injection of ketamine (4.5 mg/kg, Ketamine 1000) combined with medetomidine (45 µg/kg, Domitor). Each baboon was positioned in ventral decubitus with the head stabilised using foam supports, and a routine health check was performed. Before recordings, animals were intubated and sevoflurane (3–5%, Sevotek) was administered. Subjects were then prepared for the experiment, and sevoflurane was discontinued after installation of the EEG recording set-up. The following anaesthesia procedure depended on the subject, as described below.

### Subject-specific protocol

#### Pilot tests (Fidji and Chet)

Following common veterinary guidelines, propofol (Propovet®) was administered intravenously at 0.2 mg/kg/h following sevoflurane termination (Table 1), throughout the rest of the experiment. Ten minutes after propofol administration, we conducted a 2-minute auditory ERP experiment and a 17-minute auditory rhythmic discrimination task (not included in this manuscript).

#### Follow-up experiment (Ozone)

Because results (see below) from the previous two subjects suggested possible subject-dependent dosage effects, we aimed to better characterise this phenomenon. Thus, for the third subject, propofol was initiated at the lower dose of 0.15 mg/kg/h immediately following sevoflurane termination. After maintaining this rate for 5 minutes, the dose was reduced to 0.10 mg/kg/h for approximately 1 hour, during which an auditory ERP experiment and an auditory rhythmic discrimination task were conducted. Following this period, the propofol infusion rate was increased to 0.20 mg/kg/h for 5 minutes and finally elevated to 0.25 mg/kg/h for the last 5 minutes (Table 1).

### EEG recording

All three subjects were similarly prepared for the EEG recording. First, the temporal areas were shaved. The scalps were then washed with soap and a scrubbing skin gel (Neurprep, Spes Medica), and then dried with paper towels. Eight gold cup electrodes were positioned on the scalp at locations Cz, Fz, Pz, Oz, C3, T3, C4, and T4, along with a reference electrode at FCz and a ground electrode on the nose (Supp fig. 1). The electrodes were secured using conductive paste (SAC2, Spes Medica) and medical tape. They were then connected to a G.Nautilus amplifier (g.tec medical engineering GmbH, Austria), positioned on the operating table beside the baboons, with data wirelessly transmitted to a receiver connected to a DELL recording laptop. Electrode impedance was maintained below 30 kΩ, and data were recorded at 500 Hz sampling rate. After securing the electrodes, a wired headphone (JVC HA-S500) connected to the stimulus computer was placed on the ears of the anaesthetised baboon. Ten minutes after the start of the propofol injection, the auditory stimuli (i.e., the ERP or rhythm tasks) were broadcast to the baboon through the positioned headset.

### EEG data

All EEG preprocessing steps were done in MATLAB using the Fieldtrip toolbox and custom-written scripts. EEG data were bandpass filtered between 0.1 and 150 Hz (order =3) and a DFT filter was applied at 50, 100, and 150 Hz, as well as a third-order bandstop filter between 48-52Hz to remove background electrical noise. (Table 1).

#### ERP analysis

EEG data were bandpass filtered between 0.5 and 30Hz. Recordings were epoched from 0.5 s pre-stimulus onset to 1s post-stimulus onset. Artefact rejection was done through an amplitude-based rejection fieldtrip function, with a cut-off value at ± 100 *mV*. A final visual inspection of all trials was used to remove any remaining noisy trial that escaped the rejection procedure. In the case of Subject Ozone, the C3 electrode was shown to contain more than 80% of the artefacts and was removed from further analysis. For all subjects, EEG data were segmented and aligned to the onset of each event using the Fieldtrip Ft_timelockanalysis function^26^, then averaged across trials and electrodes, to produce an event-related potential (ERP). To normalize the data and allow comparisons across individuals, a z-score transformation was applied.

#### Sleep scoring

EEG signals were band-pass filtered between 0.3 and 30 Hz and segmented into 30-second epochs. Sleep scoring was performed offline by two experienced scorers (V.S. and C.L.), who were blind to the experimental conditions. To better assess variations in sleep depth in response to different anaesthetics and doses, we used an adapted version of the former 4-stage Rechtschaffen and Kales (1968) classification (instead of the current 3-stage AASM classification)^27^. In our system, stage N3 was defined as the presence of delta waves exceeding 50% of the epoch, while stage N4 was identified by the predominance of large amplitude delta waves throughout the entire 30-second epoch. The interrater agreement was moderate (Cohen’s κ coefficient = 0.44), and any discrepancies between the two scorers were resolved by jointly reviewing the data and reaching a consensus. Central tendency (mode) was then computed for each subject and condition to identify the dominant sleep stage under respective anaesthetic types and doses.

### Software

All analyses and visualisation were done using Matlab version 2023a and R version 4.1.2 (2021-11-01) with the packages ggplot2^28^, and tydiverse ^29^.

## RESULTS

We first conducted ERP recordings on two anaesthetised female baboons, aged 12 years (Fidji) and 16 years (Chet), while monitoring their sleep stages. The animals were maintained under propofol anaesthesia at 0.2 mg/kg/h, aiming to induce a light sleep stage. While our first subject (Fidji) displayed a light sleep stage, with a predominance of stage 1 sleep during the experiment (fig 1a, c), our second subject (Chet), although anaesthetised with the same dose of propofol exhibited a deeper sleep stage, with a predominance of stage 3 sleep (fig 1b,c). Interestingly, and in accordance with the sleep assessment, auditory ERP responses in Fidji show a mean minimum amplitude of -1.72 at 108 ms within the 80–120 ms post-stimulus window and a mean maximum amplitude of 1.44 at 186 ms within the 150–250 ms window (fig 1d). This indicates that the classical N100 and P200 components are present under propofol anaesthesia in this subject. In contrast, ERP responses in Chet exhibit a mean minimum amplitude of -1.207 at 80 ms within the 80–120 ms window and a mean maximum amplitude of -0.21 at 0.156 ms within the 150–250 ms window (fig 1e), indicating that the classical N100 and P200 components are absent in her case.

**Figure 1.**
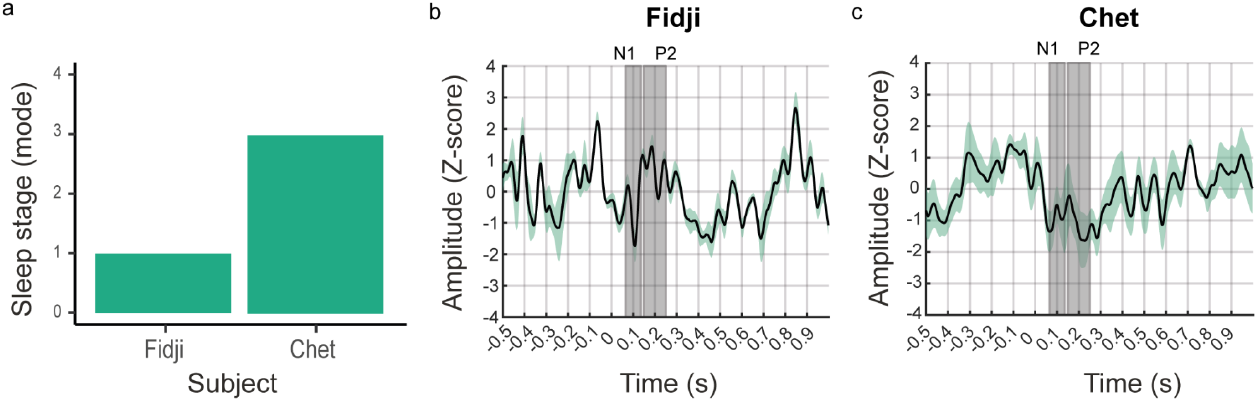
Impact of propofol infusion at 0.2mg/kg/h on vigilance states and auditory ERP (0.2 mg/kg/h). a) Central tendency sleep stage (mode) under Propofol anaesthesia in Fidji and Chet. b) Auditory ERP response (n=42) averaged across all electrodes recorded from the female baboon Fidji during propofol infusion, showing the presence of classical auditory components N100 and P200 c) Auditory ERP response (n=40) averaged across all electrodes recorded from the female baboon Chet during propofol infusion, showing the absence of classical auditory components N100 and P200.

To investigate the discrepancy in sleep and auditory responses between Fidji and Chet under propofol anaesthesia, we monitored sleep stages at varying propofol doses in a third 6-year-old female baboon (Ozone). Additionally, we recorded auditory ERP responses under sevoflurane and at a minimal propofol dose (0.10 mg/kg/h). At the start of the experiment, Ozone was under sevoflurane anaesthesia, exhibiting deep sleep stages (Central tendency = stage 4). Upon transitioning to propofol at 0.15 mg/kg/h, she remained in deep sleep stages (NREM stage 4). The first signs of sleep lightening appeared 6 minutes after the switch, becoming more pronounced 15 minutes post-transition, following propofol decrease at 0.10 mg/kg/h. During the auditory experiment, Ozone alternated between light and deep sleep stages, with light sleep predominating (75%). At the end of the experiment, following an increase in the propofol infusion rate, deep sleep stages reappeared 5 minutes after the adjustment, with a complete transition to deep sleep occurring 9 minutes later (t+14minutes;fig 2 a, 2b).

**Figure 2:**
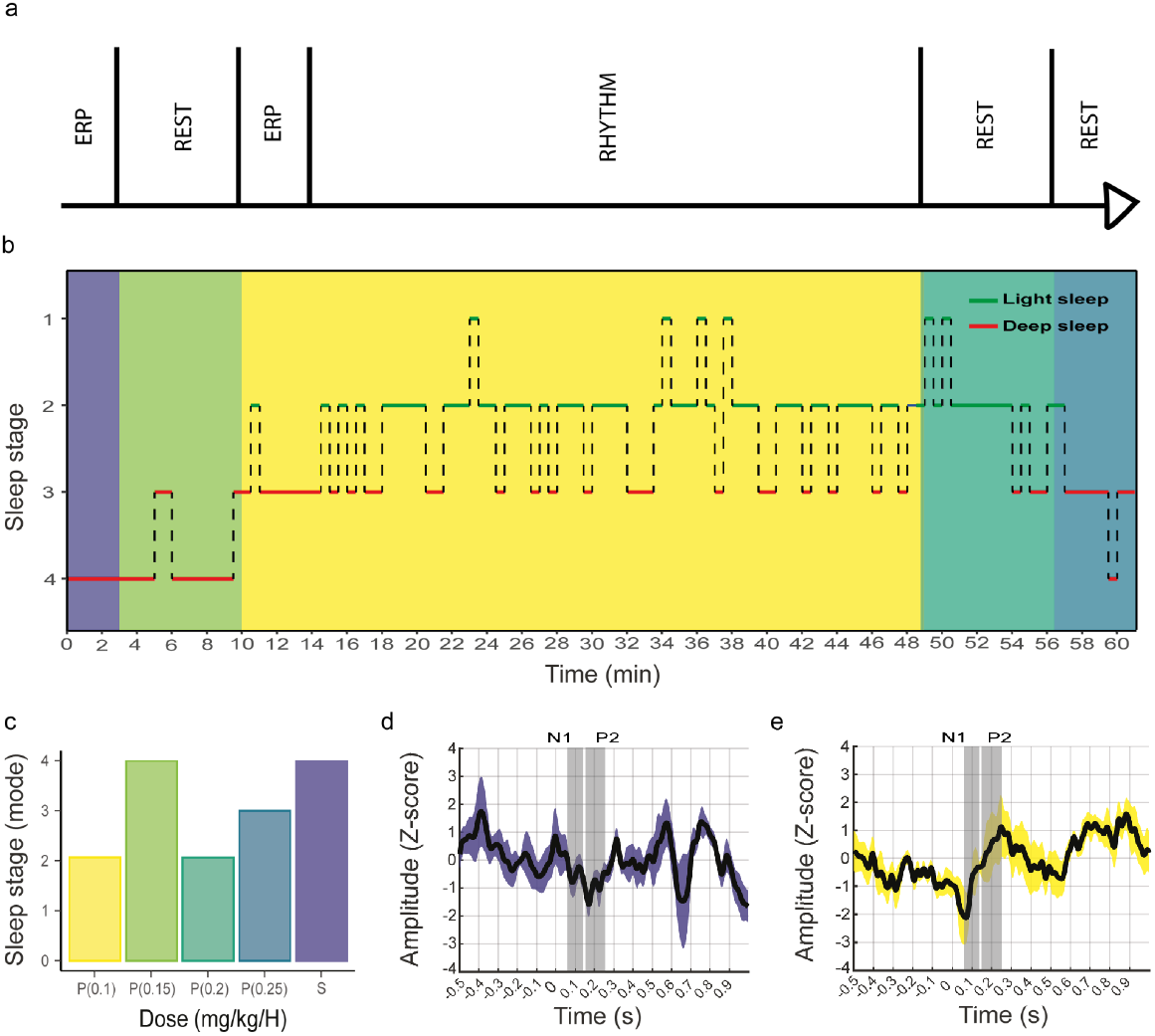
Impact of propofol infusion rate on vigilance states and auditory ERP. a) Experiment outline. b) Hypnogram showing the evolution of sleep stages across time under different Sevoflurane and Propofol infusion rates. c) Central tendency sleep stage (mode) under Sevoflurane and Propofol at different infusion rates. d) Auditory ERP response (n=76) averaged across all electrodes recorded during Sevoflurane infusion, showing the absence of classical auditory components N100 and P200. e) Auditory ERP responses (n=76) averaged across all electrodes recorded during Propofol infusion at 0.1mg/kg/h, showing the presence of classical auditory components N100 and P200.

The ERP responses under Sevoflurane, show a mean minimum amplitude of -0.80 at 86 ms within the 80–120 ms post-stimulus window and a mean maximum amplitude of -0.44 at 250 ms within the 150–250 ms window. These findings indicate that Sevoflurane anaesthesia abolished the classical N100 and P200 components. In contrast, ERP responses under light propofol anaesthesia exhibited a mean minimum amplitude of -1.87 at 80 ms within the 80–120 ms window and a mean maximum amplitude of 1.16 at 248 ms within the 150–250 ms window, confirming that light propofol anaesthesia preserves the classical N100 and P200 components.

## DISCUSSION

Performing EEG in non-human primates presents significant challenges due to motion artefacts, the fragility of recording equipment, and ethical considerations. Despite these obstacles, EEG remains crucial for advancing our understanding of brain mechanisms, particularly regarding the evolution of cross- and conspecific vocalisation processing. In light of these challenges, we aimed to assess the feasibility of using propofol anaesthesia as an alternative approach to minimise motion artefacts and stress, while enabling high-quality, non-invasive EEG recordings. By examining the effects of propofol on sleep depth and auditory processing, we sought to determine its suitability for conducting passive EEG experiments in NHPs.

Discrepancies in sleep levels and auditory responses between Fidji and Chet, despite being anaesthetised with the same dose of propofol at an injection rate of 0.2 mg/kg/h, highlight significant individual variability in propofol response (fig 1). This variability is unlikely to be related to body weight, as the injection rate was adjusted accordingly. However, since propofol is lipophilic, body composition, a factor not typically measured in NHPs, could have influenced the anaesthetic response. Additionally, factors such as the animals’ medical histories and the number of previous anaesthetic procedures they have undergone, which were not available to us, might also contribute to these differences. Consequently, such variability underscores the importance of starting with the lowest recommended dose of propofol anaesthesia, currently set at 0.1 mg/kg/h according to veterinary guidelines, to assess the subject’s response and adjust the dosage as needed.

In the third subject Ozone, sleep levels under propofol anaesthesia exhibited a dose-dependent response, with deeper sleep observed at an infusion rate of 0.25 mg/kg/h compared to 0.1 mg/kg/h (fig. 2c). While propofol at 0.1 mg/kg/h predominantly triggers light sleep stages, an alternation between NREM2 and NREM3 still occurs. This suggests that further reducing the dose could potentially favour an alternation between lighter sleep stages, best suited for exploring auditory processing. Continuous monitoring of sleep levels throughout the experiment also revealed a residual effect of Sevoflurane anaesthesia. Sleep levels started to decrease 6 minutes after the start of propofol and stabilised around 15 minutes post-initiation, emphasising the need for a waiting period between anaesthesia transitions and the start of auditory experiments. Similarly, when the propofol infusion rate was increased from 0.1 mg/kg/h to 0.2 mg/kg/h, changes in sleep stages took about 5 minutes to manifest (fig. 2a, 2b).

Importantly, in our two subjects exhibiting predominantly light sleep stages during the auditory ERP protocol (Fidji and Ozone), brain signals revealed the presence of the classical auditory components N100 and P200, indicating that basic auditory processing is preserved under light propofol anesthesia (Fig. 1d, 2e). This finding highlights the feasibility of using light propofol anesthesia for non-invasive brain recordings in non-human animals, preserving essential neural functions while limiting the need for invasive methods.

Performed alongside routine veterinary care, light propofol anaesthesia for brain recording in non-human animals offers significant practical benefits from both ethical and experimental perspectives. The key advantages of this protocol are: 1) stress reduction compared with traditional restraint methods for both the experimenter and the animal subject, 2) the naturalness of the possible experimentations, and 3) the amount of data that can be acquired as more animals can be tested without harm. Given that propofol does not inhibit pulmonary function, it is also widely used in human medicine for anaesthesia without the need for artificial ventilation ^30,31^. Therefore, an adapted propofol protocol could potentially be applied to wild, untamed animals, facilitating EEG recordings in natural and uncontrolled environments under veterinary supervision. While our protocol involved shaving the scalp to improve electrode contact, it is worth noting that similar studies in humans have achieved high-quality data with the same electrodes on unshaved subjects^32^, suggesting the potential of minimising the invasiveness of the procedure in future experiments. Alternative methods, such as the use of spider electrodes or other innovative electrode types, could also be explored to minimise the procedure’s invasiveness and enhance its applicability in field settings.

Overall, our findings validate the use of light propofol anaesthesia for auditory-related tasks in non-human animals, while highlighting key considerations for its use. Starting with a low dose of 0.1 mg/kg/h or even lower, and adjusting the dose based on individual responses is essential to preserve basic auditory processing in NHP’s brain. A waiting period of 15 minutes should be observed between anaesthesia initiation and the start of the experiment to allow sleep to stabilise. Any increase in dose or adjustment in infusion rate should be closely monitored, as changes in sleep stages may take several minutes to appear, significantly affecting auditory processing. This approach not only provides an ethical and practical solution for conducting brain recordings but also opens the door for future research that could explore more complex auditory processing, such as mismatch negativity (MMN) in an auditory oddball paradigm, local-global paradigms^33^, as well as long natural processing experiments. This would help assess whether higher-order auditory functions are also preserved under light propofol anaesthesia, paving the way for more advanced investigations into neural processes in both captive and wild non-human animals.

## ETHICAL STATEMENT

All animal procedures were approved by the “C2EA−71 Ethical Committee of neurosciences” (INT Marseille) under the number APAFIS##46189-2023111410456237 v5, and have been conducted at the Station de Primatologie under the number agreement G130877 for conducting experiments on vertebrate animals (Rousset-Sur-Arc, France). All methods were performed in accordance with the relevant French law, CNRS guidelines and the European Union regulations (Directive 2010/63/EU).

## ACKNOWLEDGEMENTS

The NCCR Evolving Language, Swiss National Science Foundation Agreement Nr. #51NF40_180888, funded this work. We are very grateful to the Station de Primatologie CNRS, particularly the animal care staff and technicians, for their critical involvement in this project. A.M has received funding from the European Research Council under the European Union’s Horizon 2020 research and innovation programme grant agreement No 716931—GESTIMAGE—ERC-2016-STG, as well as French National Research Agency ANR-23-CE28-0029-01 (BABONTO), ANR-16-CONV-0002 (ILCB).

## CONTRIBUTIONS

T.P, E.D, D.G, A.M and A-L.G conceptualised the project. T.P, M.S, M.I, V.M adapted and refined the anaesthetic protocol. M,I and V.M performed animal handling and anaesthesia. T.P, M.S, A.M and D.G conducted EEG data collection. T.P performed EEG data analysis and visualisation. C.L and V.S performed sleep scoring. T.P wrote the initial draft. All authors contributed to the review and editing of the manuscript. E.D and A-L.G provided supervision. A-L.G provided resources and funding for the project.

## SUPPLEMENTARY MATERIALS

**Supp Figure 1 :** Electrodes placement on the baboon skull.

