## Supplementary Figure 1 for "Light propofol anaesthesia for non-invasive auditory EEG recording in unrestrained non-human primates"

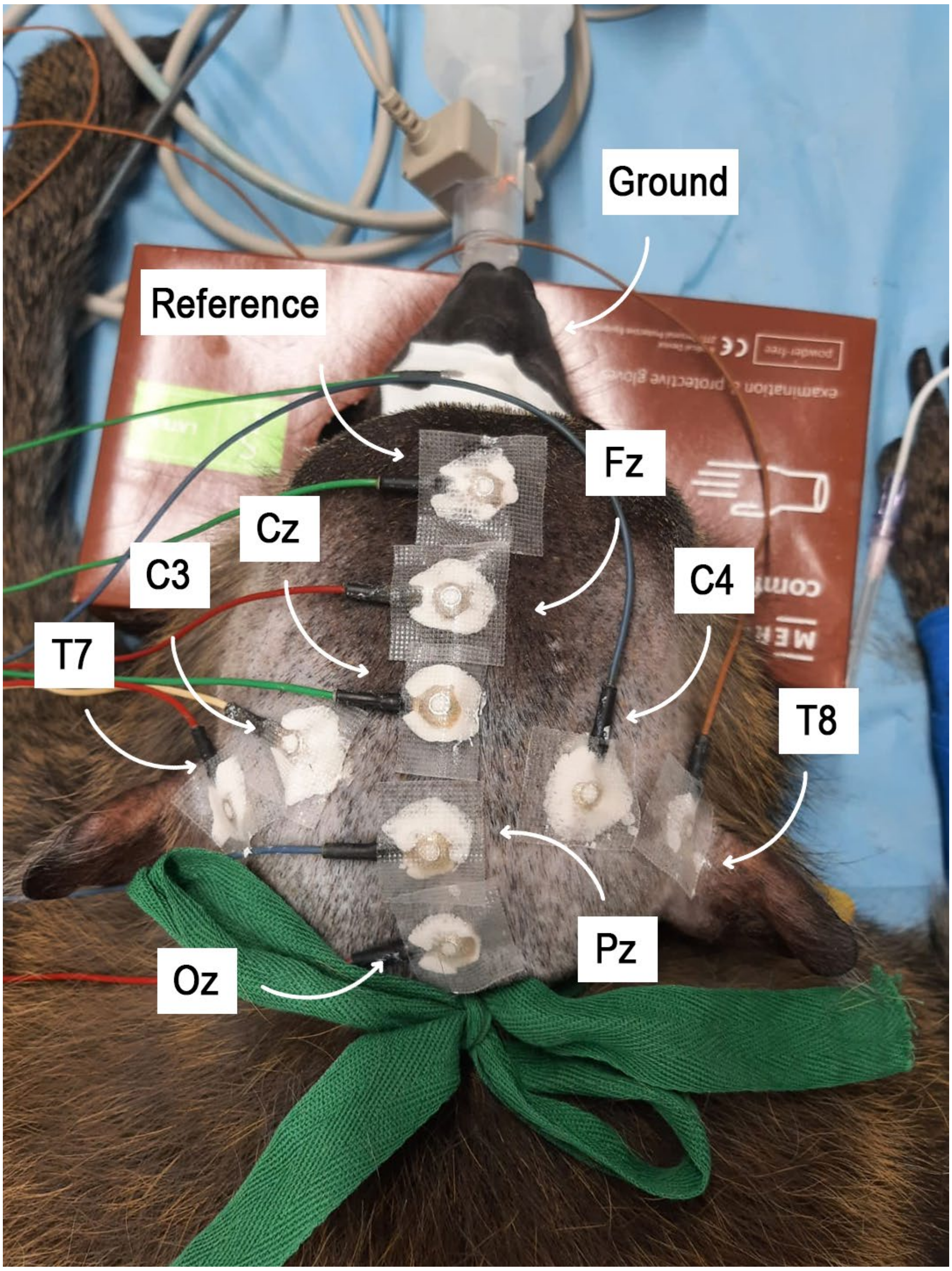


**Supplementary Figure 1 : Electrode placement.** Placement of electrodes for EEG recording on Ozone according to the 10-20 EEG layout.
